# Multi-omic profiling of pleural effusions: a reservoir-incubator for non-invasive insights into tumor heterogeneity

**DOI:** 10.64898/2026.01.27.701856

**Authors:** Paula Nieto, Ginevra Caratù, Maria Mulet, M. Alba Sorolla, Virginia Pajares, Domenica Marchese, Ana Pardessus, Rubén Osuna-Gómez, Albert Rafecas-Codern, Pere Serra-Mitjà, José M. Porcel, Silvia Vidal, Holger Heyn, Juan C. Nieto

## Abstract

Pleural effusions (PEs) are common in advanced cancer and are routinely sampled for cytological evaluation; however, their full potential as a source of tumor and immune information remains underexplored. Although malignant PEs (MPEs) are increasingly used for liquid biopsy, their cellular complex interactions have not been resolved at high resolution.

Here, we present the first comprehensive single-cell and multi-omic atlas of PEs, integrating scRNA-seq, TCR-seq and whole-exome sequencing with lung, breast, or ovarian cancer, as well as non-malignant heart failure controls. We profiled over 200,000 cells and identified more than 30 immune, stromal, and tumor subpopulations across disease contexts. The tumor cell signature in MPEs recapitulates the genomic and transcriptomic features of matched primary tumors, supporting their use for non-invasive tumor profiling. In contrast, immune landscapes varied by disease: lung cancer MPEs were enriched in exhausted CD8⁺ T cells, breast MPEs showed CD4⁺ T helper cell dominance, and ovarian and heart failure PEs harbored tissue-resident myeloid cell populations.

These data reveal that PEs are dynamic, disease-specific ecosystems shaped by tumor-driven immune remodeling. By leveraging their cellular diversity, we position MPEs as a powerful platform for studying metastatic adaptation and immune responses *in vivo*, with direct implications for diagnostics, immune monitoring, and precision oncology.

## Introduction

Pleural effusion (PE), a pathological accumulation of fluid in the pleural cavity, occurs in diverse conditions, including congestive heart failure, infections, and advanced malignancies^1^. In malignant pleural effusion (MPE), tumor cells are shed into the pleural space, most frequently in lung, breast, and ovarian cancers^2,3^. In addition to causing respiratory compromise, MPE represents a complex and poorly understood tumor microenvironment (TME), shaped by vascular leakage, immune suppression, and stromal remodeling^4^.

While PE is routinely sampled for diagnosis and symptom relief, cytological analysis often lacks sensitivity, thereby, limiting its diagnostic value^5^. Nonetheless, PE harbors abundant tumor and immune cells, as well as soluble mediators, offering a potential window into the biology of metastatic progression^6^. Recent single-cell studies have begun to reveal the immune diversity within MPEs; however, most studies have focused on individual cancer types or the immune compartment alone, leaving the full cellular complexity, and tumor–immune interactions largely unexplored^7–10^.

Here, we present an integrated high-resolution cellular atlas of malignant and non-MPEs using single-cell RNA sequencing, TCR sequencing, whole-exome sequencing, and bulk transcriptomics. Across 31 patients with lung, breast, ovarian cancer, or heart failure, we profiled over 200,000 cells spanning the tumor, immune, and stromal compartments. We identified disease-specific programs of immune infiltration, tumor heterogeneity, and T cell dysfunction. Notably, we defined a robust MPE transcriptional signature with potential diagnostic utility and showed that PE-derived tumor cells recapitulate primary tumor biology at genomic and transcriptomic levels. These findings establish PE as a minimally invasive, clinically accessible resource for dissecting the metastatic TME and guiding personalized cancer care.

## Results

### Multi-modal analyses of the pleural effusion microenvironment reveal disease-specific cellular landscapes

To comprehensively profile the PE microenvironment across malignant and non-malignant conditions, we performed scRNA-seq and paired TCR sequencing of PE samples from 31 patients with lung cancer (LC), breast cancer (BC), ovarian cancer (OC), and heart failure (HF). Additionally, we validated our findings using whole-exome sequencing (WES), and massively parallel bulk TCR sequencing in four of these patients. Clinical metadata revealed a heterogeneous cohort with a variable cytological tumor detection, and response status. The time from tumor diagnosis to PE ranged from diagnosis to over 8,000 days, reflecting the temporal diversity in metastatic progression (**Fig. 1a, Sup. Table 1**). Each sample yielded between 815 and 8,135 high-quality cells (MEAN = 5364.97, SD = 1919.75), resulting in a total of over 150,000 cells for downstream analysis (**Sup. Fig. 1a**). Prior to sequencing, we implemented a FACS-based enrichment strategy using EPCAM and CD45 antibodies to enrich two key compartments: CD45⁻*EPCAM*⁺ tumor cells and *EPCAM*⁻*CD45*⁻ double-negative cells (**Sup. Fig. 1b**). This strategy balanced the overwhelming dominance of *CD45*⁺ immune cells, which can comprise up to 95% of PE cells^11^, and enhanced our ability to increase the resolution of tumor and stromal populations (**Sup. Fig. 1c**).

We analyzed a total of 168,448 high-quality single cells from PE samples and annotated 10 major cell populations, including immune subsets (CD4+ and CD8+ T cells, B cells, NK cells, dendritic cells, monocytes and macrophages), and tumor and stromal components (tumor cells, fibroblasts and mesothelial cells) (**Fig. 1b, Sup. Fig. 1d**). In general, immune populations dominated most samples, although the proportions of each subtype varied considerably, tumor cells were detected in different proportions in LC, BC, and OC MPE but not in HF samples, consistent with their non-malignant origin (**Fig. 1c**). Differential abundance analysis comparing malignant and non-malignant effusions, revealed that MPEs displayed a reduction of resident populations including mesothelial cells, macrophages, and fibroblasts, suggesting that tumor progression disrupts normal resident cell homeostasis (**Fig. 1d**). Furthermore, compositional analysis revealed disease-specific enrichment patterns: B cells tended to increase in LC, and macrophage/monocyte compartments were reduced in most cancers (**Fig. 1e**). As our most abundant disease group, we used LC as a “reference group” for all the statistical comparisons. Additionally, comparing the CD4+/CD8+ T cell ratio across diseases showed elevated ratios in BC compared to LC, hinting at distinct immune landscapes of T cells across tumor types (**Fig. 1f**), consistent with previous findings in the primary TME^12–16^. To further investigate the immune polarization observed in PE, we annotated over 100 distinct cell populations spanning the stromal, tumor, and immune compartments, generating a comprehensive reference framework to support in silico hypothesis testing of MPE development mechanisms (**Sup. Fig. 1e**).

These findings highlight the PE as a dynamic and accessible in vivo incubator for tumor and immune cells. Its heterogeneous composition, shaped by both metastatic tumor activity and immune responses, offers a powerful window into the evolving tumor microenvironment.

**Figure 1:**
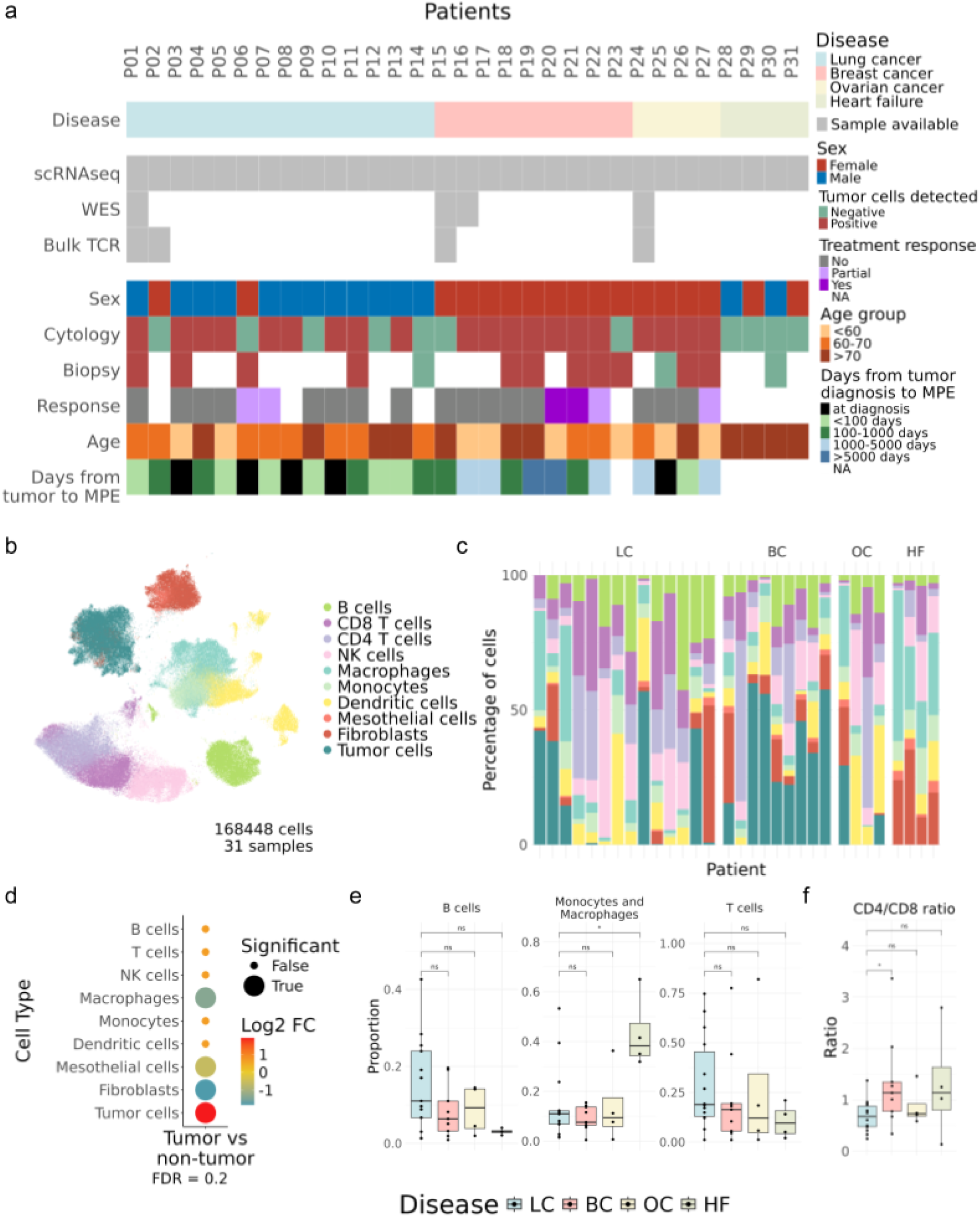
**a**, Square chart showing all PE samples and sequencing technologies used for each of them along with several clinical and demographic parameters including sex, age at diagnosis, treatment response, detection or lack thereof of tumor cells through cytology or biopsy and days from tumor diagnosis to MPE. **b**, Integrated UMAP of all cells sequenced by scRNA-seq in PE across patients and diseases (168448 cells), colored by cell type. **c**, Bar plot showing the PE composition of all patients at the general level of resolution. Cell type color scale is the same as in b. **d**, Dot plot showing the results of applying scCODA on the general annotation comparing all the tumor groups against the heart failure. The false-discovery rate was set at 0.2. **e**, Box plots showing the different cell type proportions per patient and disease. The Wilcoxon rank-sum test was used to compare all disease groups to the lung cancer group. **f**, Box plots showing CD4+ to CD8+ T cell ratio per patient and disease. A Wilcoxon rank sum test was applied to compare all disease groups against the lung cancer group.

### Tumor cell programs reveal heterogeneous adaptation across cancer types

To dissect the cellular diversity within the non-immune compartment of PE, we analyzed over 48,000 *CD45*⁻ cells in MPE, revealing a heterogeneous landscape of tumor (*EPCAM*⁺) and stromal phenotypes. We identified twelve transcriptionally distinct populations, including non-malignant mesothelial and stromal-like subsets, myofibroblasts, EMT-like (Epithelial-to-Mesenchymal Transition) cells, and multiple tumor states (**Fig. 2a, Sup. Fig. 2a,b**). The presence and abundance of these states varied markedly across patients and cancer types, highlighting pleural effusions as highly dynamic and patient-specific niches for tumor cell evolution (**Sup. Fig. 2b,c**). Statistical differential abundance testing using LC as the reference group, owing to its higher sample number, cytological diversity and clinical localization, revealed disease-specific enrichment patterns: SCGB1D2+ and MUC+ tumor cells, among others, were enriched in BC, whereas OC was enriched in stromal-like tumor cells (**Fig. 2b**). Unsurprisingly, HF showed a depletion of all tumor subgroups and an increase in resident fibroblast populations.

Using CancerSEA’s^17^ single-cell tumor profile signatures, we further explored the predominant functional programs of tumor cells across diseases. This revealed higher EMT, angiogenesis, and stemness signatures in OC-derived effusions, and increased cell cycle and apoptotic activity in BC (**Fig. 2c**). This variability reinforces the notion of the PE as a human incubator, where tumor cells adopt distinct adaptive strategies depending on their tissue of origin and local selective pressures.

To assess the extent to which these tumor subtypes mirror primary tumors, we compared our data with TCGA samples, including between 500 and 1000 bulk transcriptomes per disease (**Sup. Fig. 2d**). Tumor cell states identified in PE, including immunogenic, proliferative, and stromal-like subtypes, showed high similarity to transcriptomic patterns in lung adenocarcinoma (LUAD), breast invasive carcinoma (BRCA), and cervical squamous cell carcinoma and endocervical adenocarcinoma (CESC), suggesting that pleural effusions recapitulate key aspects of the primary tumor microenvironment (**Fig. 2d**). Cross-validation with scRNA-seq cell signatures from recent primary tumor studies and naive disease enrichment analyses further confirmed the conservation of primary tumor identities (**Fig. 2e, Sup. Fig. 2e**). Finally, combined copy number variation analysis (CNV) from WES and scRNA-seq confirmed the tumor identity of malignant cells and aligned with the inferred heterogeneity but suggested tumor detection through MPE (**Sup. Fig. 3**). Together, these findings show that PEs are not passive fluid collections but active ecological niches that preserve the transcriptional plasticity of tumor cells. Their study enables a detailed resolution of context-specific metastatic programs with diagnostic and therapeutic relevance.

**Figure 2:**
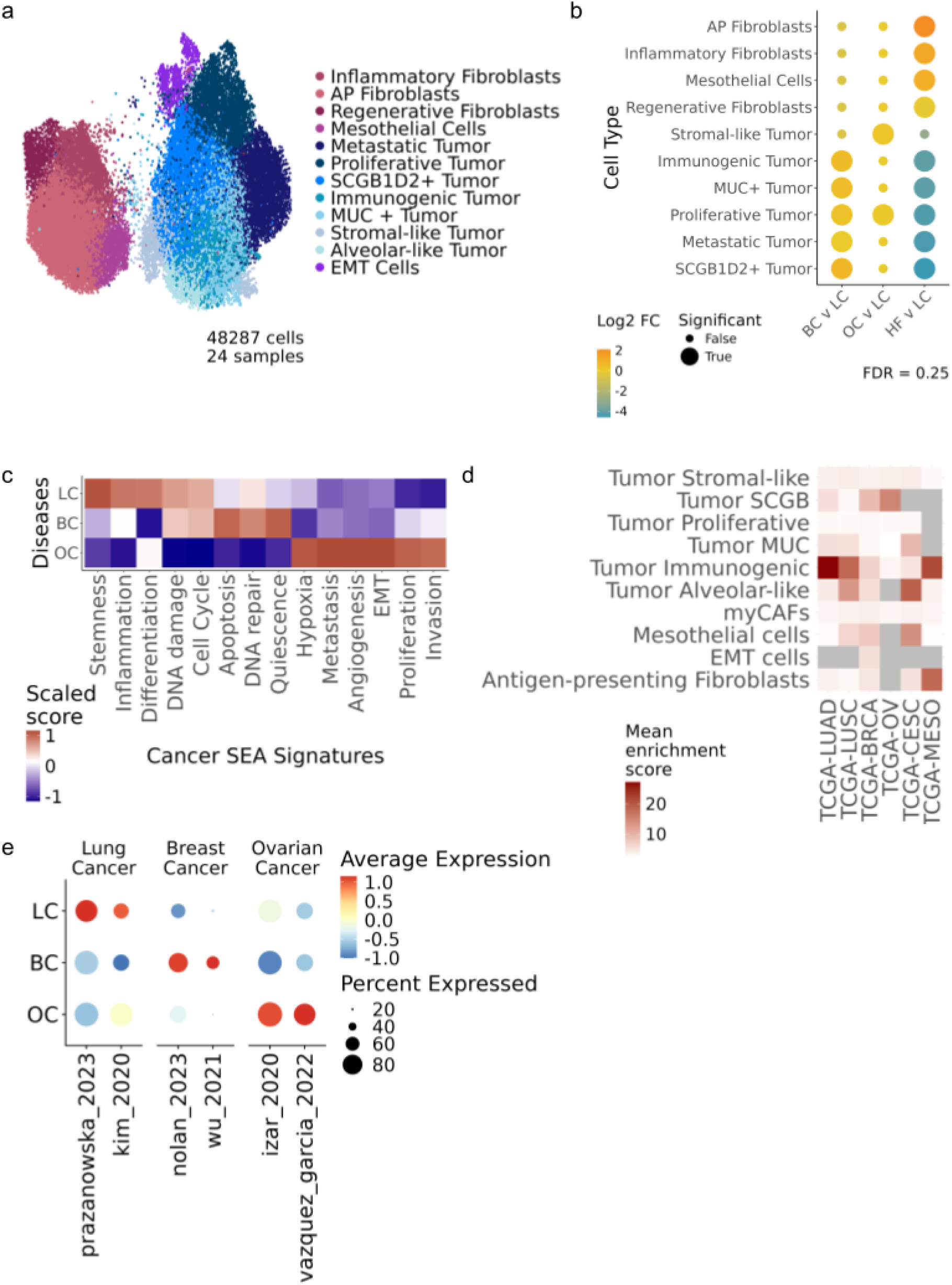
**a**, Integrated UMAP of all *CD45-* cells sequenced by scRNA-seq in PE across all patients and diseases (48287 cells), colored according to cell type. **b**, Dot plot showing the results of applying scCODA on *CD45-* cells comparing all disease groups against heart failure. The false-discovery rate was set at 0.3. **c**, Heatmap showing the different signature scores in lung, breast and ovarian cancer cells only. Scores are scaled. **d**, Heatmap showing the mean cell type enrichment score per TCGA project (cancer type). BRCA, breast invasive carcinoma; LUAD, lung adenocarcinoma; LUSC, lung squamous cell carcinoma; OV, ovarian serous cystadenocarcinoma; CESC, cervical squamous cell carcinoma and endocervical adenocarcinoma; MESO, mesothelioma. **e**, Dot plot showing the average expression and percentage of cells expressing public cancer-type-specific signatures in PE cancer cells.

### A diagnostic gene signature detects tumor presence in pleural effusions, improving malignancy detection

After validating and characterizing the tumor compartment as a window into the metastatic environment, we developed a transcriptional fingerprint of MPE to enhance tumor cell detection within the PE. This molecular tool addresses the limited sensitivity of cytology in traditional clinical practices, enabling more accurate diagnosis of neoplasias. We performed differential expression analysis between *CD45*⁻ cells from MPE and HF PE groups. This yielded a robust MPE signature composed of genes such as *CEACAM5, MUC1, CLDN4*, and *SCGB2A2*, which are known markers of epithelial transformation and malignancy (**Fig. 3a**).

The signature was consistently enriched only in tumor cells from pleural effusions associated with LC, BC and OC, but not in non-malignant cell types (**Fig. 3b**). Next, we evaluated the diagnostic power of the MPE signature across primary tumors to confirm the specificity of tumor cell detection. We scored the signature in public scRNA-seq datasets from matched primary tumors from Salcher et al. 2022, Wu et al. 2021 and Vázquez-García et al. 2022 (see **Methods**). This validated the specificity of our signature in identifying tumor cells (**Fig. 3c**). The score positively correlated with the proportion of tumor cells present in each sample (R = 0.78, p = 2.9e-07), confirming its specificity and sensitivity in detecting malignant content (**Fig. 3d**). Importantly, when compared to clinical pathology assessments, MPE signature scores distinguished cytology- or biopsy-positive from negative cases (**Fig. 3e**), demonstrating its potential to improve the detection of tumor cells in PE compared to standard diagnostic cytology, a highly relevant feature, given the pressing clinical need to improve traditional detection methods across tumor entities^18^.

To evaluate its relevance beyond our cohort, we scored our MPE signature onto bulk RNA-seq primary tumor samples from TCGA. The MPE signature was broadly enriched in lung adenocarcinoma (LUAD), breast invasive carcinoma (BRCA), and ovarian serous cystadenocarcinoma (OV) samples but was absent in mesothelioma (MESO), suggesting that it captures canonical epithelial malignancy features rather than generic pleural transcriptional programs (**Fig. 3f**). Finally, to benchmark the detection sensitivity of our signature, we performed bulk RNA-seq of LC MPEs and non-MPEs (HF) with different tumor cell numbers quantified by flow cytometry. We observed a positive correlation between the MPE score and the absolute number of tumor cells recovered per sample (**Fig. 3g**), supporting its usefulness as a quantitative proxy for malignant burden. Collectively, these results position the MPE signature as a robust and specific readout with high translational potential for the molecular diagnosis of PE.

**Figure 3:**
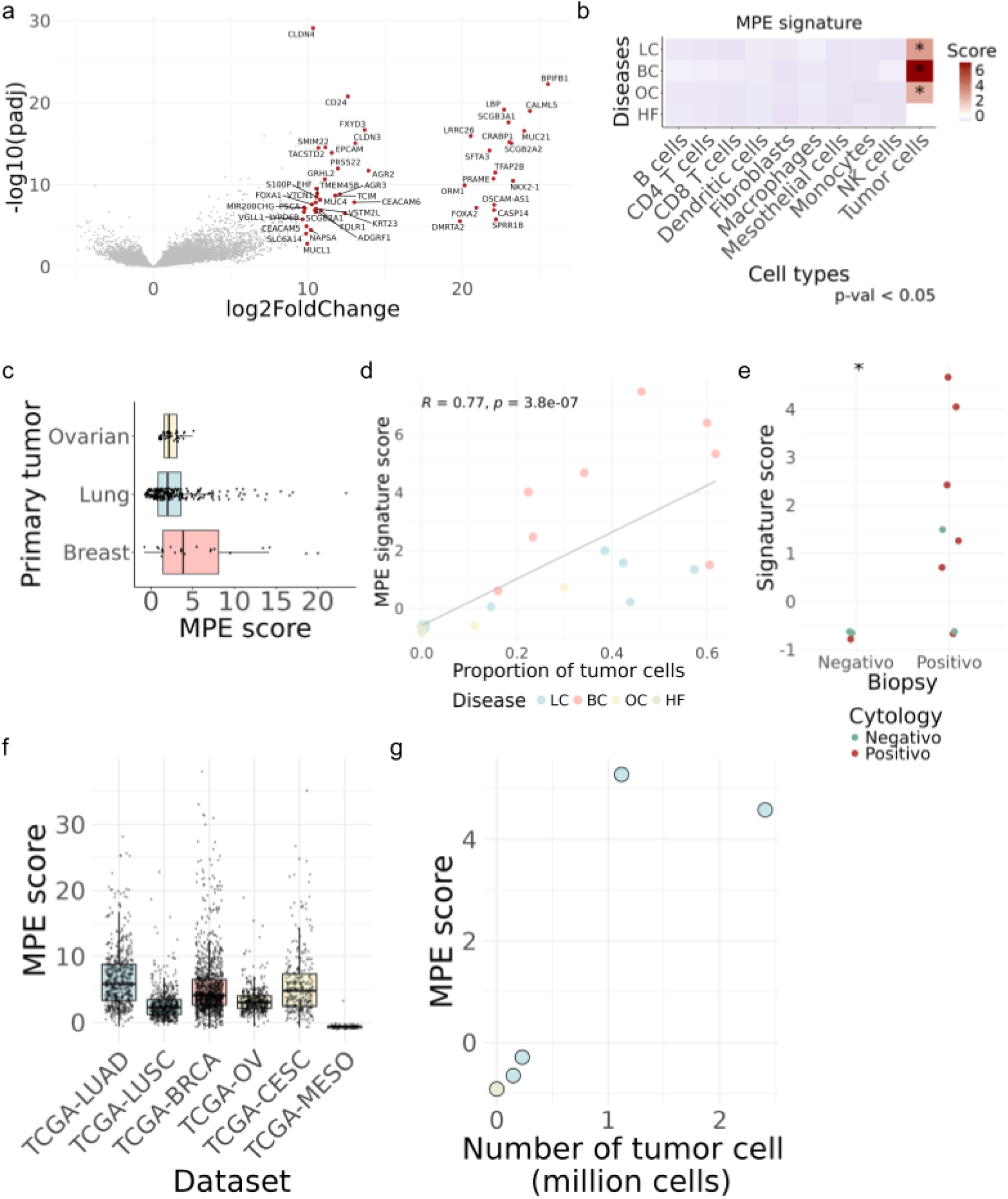
**a**, Volcano plot showing differentially expressed genes between *CD45-* cells from MPE and non-malignant PE groups. Labelled genes are the top 50 genes with higher log2-fold-change, which form our MPE signature. **b**, Heatmap showing the expression scores of the MPE signature across all cell types and disease states. Asterisks highlight scores with p-values < 0.05. **c**, MPE signature score in tumor cells from primary tumors in public scRNA-seq datasets. **d**, Correlation between the MPE signature score and the proportion of tumor cells per sample. **e**, Scatter plot showing MPE signature score across patients who tested negative/positive for the presence of tumor cells through biopsy analysis colored by the same result for the cytology test of the PE. **f**, MPE signature expression score across bulk RNA-seq samples from several TCGA projects (cancer types). BRCA, breast invasive carcinoma; LUAD, lung adenocarcinoma; LUSC, lung squamous cell carcinoma; OV, ovarian serous cystadenocarcinoma; CESC, cervical squamous cell carcinoma and endocervical adenocarcinoma; MESO, mesothelioma. **g**, Scatter plot showing MPE signature score across patients from the bulk RNA-seq validation cohort and the total initial number of tumor cells per sample (in millions).

### TCR expansion of OXPHOS CD8⁺ T cells in MPE is a predictor of treatment resistance

Having characterized the cellular landscape of the PE and identified a robust MPE tumor signature among the non-immune cells, we next moved to T cells, key effectors of anti-tumor immunity, to investigate whether their clonal architecture and transcriptional states were associated with the clinical response. TCR repertoire analysis across diseases (LC BC, OC and HF) did not reveal significant differences in clonal diversity or expansion in CD8+ T cells (**Sup. Fig. 4a**) but exhibited significantly higher clonal diversity and diminished clonal expansion of CD4+ T cells in BC than LC, highlighting the relevance of CD4+ T cells in BC compared to LC (**Fig. 4a**). However, when stratified by treatment response, patients who did not respond to treatment displayed markedly reduced TCR diversity and increased clonal expansion in their CD8⁺ T cells (**Fig. 4b**), suggestive of oligoclonal expansion associated with a dysfunctional state. However, these differences in treatment responses were not observed among CD4+ T cells (**Sup. Fig. 4b**).

We then profiled the phenotypic landscape of CD8⁺ T cells to further investigate these differences. We characterized the heterogeneity of CD8⁺ T cell composition, which ranged from naive, effector-cytotoxic, memory and terminally exhausted populations (**Sup. Fig. 4c,d,e**) with notable shifts between the responders and non-responders (**Fig. 4c**). Specifically, responders exhibited significantly increased proportions of central memory (CM) CD8⁺ T cells and a depletion of effector memory (EM), resident and exhausted CD8⁺ T cells, compared to non-responders (**Fig. 4d**). This distribution suggests that T cells in responders retain a more functional, infiltrative and less differentiated state, possibly maintaining their activation and cytotoxic potential, while exhaustion seems to play a key role in the lack of treatment response.

Next, we explored whether particular functional programs could explain these phenotypic differences. Using pathway activity scores derived from the MSigDB cancer hallmarks database^19^, we found that the oxidative phosphorylation program was significantly enriched in the cell types depleted in responder patients, particularly within proliferative, exhausted, and effector-like CD8⁺ subsets (**Fig. 4e**). These findings are consistent with previous reports linking high oxidative phosphorylation activity to T cell dysfunction and treatment failure in tumors^20,21^.

Interestingly, although the overall exhaustion scores were not statistically different between the groups (**Sup. Fig. 4f**), we observed a significant positive correlation between exhaustion and a previously defined tumor-infiltrating lymphocyte (TIL) signature from Oliveira et al. 2021, suggesting that TILs captured in PE also exhibit an exhausted phenotype (**Fig. 4f**).

We extended our analysis to four matched primary tumors (two LC, one BC and one OC; see **Sup. Table 1**) to investigate the TCR repertoire relationships between tumor-infiltrating T cells and those found in the MPE. By integrating massively parallel bulk TCR sequencing (osTCR) from primary tumors with scTCR-seq data from PE, we identified shared TCR clones between the tumor and PE compartments, highlighting their trafficking. Notably, the clones shared between the tumor and MPE were enriched for CD8⁺ effector cells, suggesting that the MPE reflects a memory-like compartment seeded by the primary tumor immune environment. The clone size in both compartments did not exhibit any correlation, implying that clonal expansion can occur independently within the compartments (**Fig. 4g**). This finding underscores the complex and dynamic relationship between T cell exhaustion, metabolic state, and therapeutic outcomes in the MPE tumor microenvironment.

Together, these data suggest that treatment resistance could be associated with a clonally expanded, metabolically rewired CD8⁺ T cell compartment in MPEs, and that metabolic profiling of T cells and their TCRs may serve as a predictor of treatment response.

**Figure 4:**
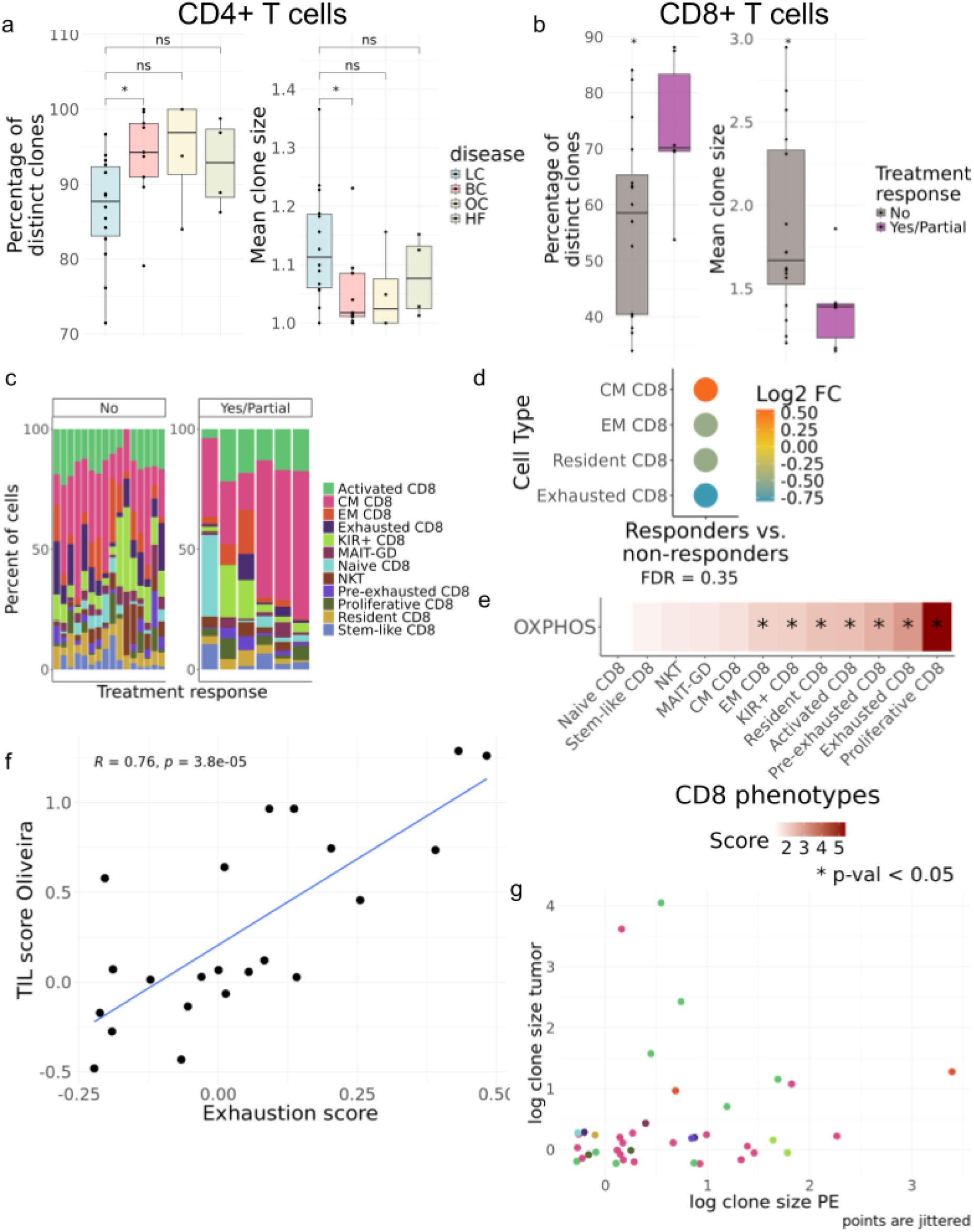
**a**, Box plots showing the percentage of distinct clones and mean clone size of CD4*+* T cells per patient and disease. The Wilcoxon rank-sum test was used to compare all disease groups with the lung cancer group. **b**, Box plots showing the percentage of distinct clones and mean clone size of CD8+ T cells per patient and treatment response. The Wilcoxon rank-sum test was used to compare the two groups. **c**, Bar plot showing CD8+ T cell composition across patients, grouped by responders and non-responders and colored according to cell type. **d**, Dot plot showing the results of applying scCODA on CD8+ T cells comparing responders and non-responders. Only populations exhibiting significant changes are shown. The false discovery rate was set at 0.35. **e**, Heatmap showing the OXPHOS signature score across CD8+ T cell phenotypes. **f**, Scatter plot showing the correlation between TIL score and exhaustion score in CD8+ T cell phenotypes grouped by treatment response. **g**, Scatterplot showing the distribution of clone sizes in logarithmic scale for those clones shared between tumor and PE. Dots are colored by the most abundant phenotype for each clone in PE. Same color scale as in **c**.

### Myeloid landscape of PE reveals disease-specific patterns of immune infiltration

Unsupervised clustering identified a diverse array of myeloid populations, including classical and non-classical monocytes, multiple dendritic cell (DC) subsets (DC1, DC2, pDC, mregDC and CD207+ DC), and tumor-associated macrophage (TAM) phenotypes such as alveolar TAM or metabolic lipid-associated macrophages (**Fig. 5a, Sup. Fig. 6a**). Differential abundance analysis revealed disease-specific shifts in the myeloid compartment when comparing LC with other conditions. BC samples were enriched for classical monocytes, suggesting a more infiltrative and potentially immunoreactive myeloid environment. In contrast, OC exhibited a depletion of many myeloid populations, including interstitial IM1 macrophages, metabolic-LA macrophages, TAM-like alveolar macrophages, classical monocytes and MDSCs. In contrast, HF was characterized by increased proportions of resident and inflammatory populations, including tissue macrophages, inflammatory monocytes, and interstitial IM1 macrophages, as well as a significant reduction of pDCs, consistent with a more localized or immunosuppressive myeloid landscape (**Fig. 5b, Sup. Fig. 6b**).

To better resolve functional specialization within the myeloid compartment, we scored individual macrophage and monocyte subpopulations for the expression of key gene tumor-associated-state signatures, including blood-derived monocytes (*BDM_TAM*), resident-like (*Res_TAM*), interstitial-like macrophages (*Interst_TAM*), pro-fibrotic (*Profibro_TAM*), pro-inflammatory (*PRO_INFL*) and anti-inflammatory (*ANTI_INFL*), tumor-associated programs. Alveolar TAMs and metabolic LA-TAMs exhibited consistently high expression across nearly all TAM-related signatures, suggesting that these populations are central hubs of macrophage plasticity and functional reprogramming within the pleural tumor microenvironment. Interstitial IM1 macrophages and inflammatory monocytes also showed elevated expression of interstitial and pro-inflammatory programs, reflecting a more activated, inflammatory state. In contrast, tissue macrophages displayed low expression across all TAM programs, with a marked reduction in anti-inflammatory and profibrotic scores, suggesting a functionally distinct and possibly homeostatic phenotype. Proliferative macrophages showed low to intermediate expression of TAM programs, whereas anti-inflammatory macrophages were enriched specifically in ANTI_INFL and Profibro_TAM signatures. This dot plot highlights how TAM-related programs segregate across macrophage subtypes, with alveolar and metabolically reprogrammed macrophages exhibiting the highest diversity and intensity of tumor-supportive gene expression (**Fig. 5c**). We then compared these myeloid phenotypes in PE with primary tumor data from the literature (Salcher et al. 2022, Wu et al. 2021 and Vázquez-García et al. 2022). Quantification of functional signature scores revealed similarities between the primary tumor environment and PE. For example, ANTI_INFL, Profibro_TAM and BDM_TAM levels were similarly conserved in LC patients. However, only similar levels of PRO_INFL and BDM_TAM were observed in BC. The ovarian primary tumor was similar to MPE OC in ANTI INFL, PRO_IFL and Profibro TAM depletion (**Fig. 5d**). Overall, the myeloid landscape in malignant pleural effusions is shaped by conserved TAM programs with some disease-specific modulation, underscoring their role in local immune remodeling and potential relevance for immunotherapeutic targeting.

**Figure 5:**
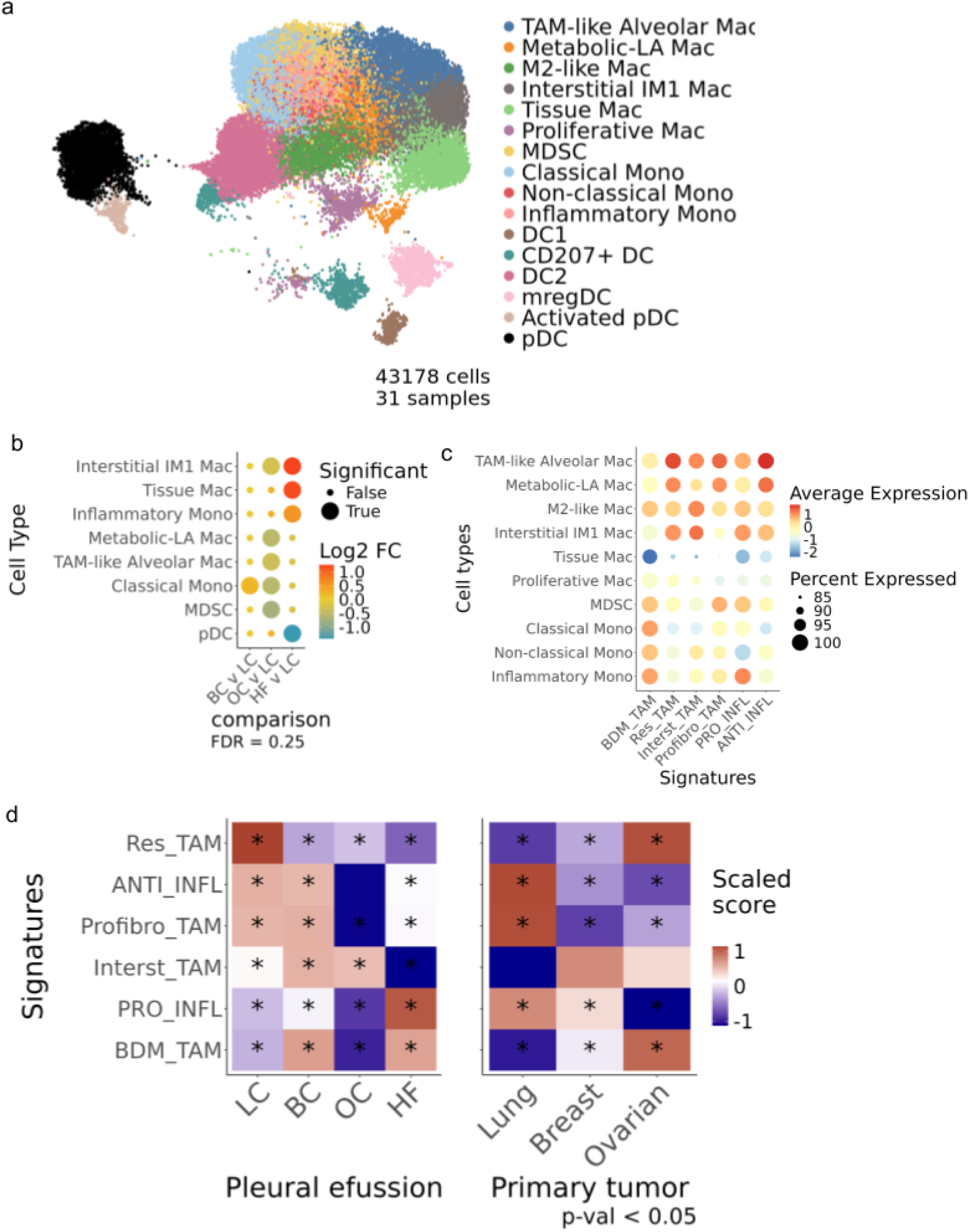
**a**, Integrated UMAP of all myeloid cells sequenced by scRNA-seq in PE across all patients and diseases (43178 cells), colored according to cell type. **b**, Dot plot showing the results of applying scCODA to the myeloid cells, comparing each disease group with lung cancer. Only the cell groups exhibiting significant changes are shown. The false discovery rate was set at 0.3. **c**, Dot plot showing average expression and percentage of cells expressing each of the selected myeloid cell signatures across all macrophage and monocyte populations **d**, Heatmaps showing pseudobulk scaled scores across myeloid cell populations in PEs (left) and primary tumors (right).

## Discussion

In this study, we present the most comprehensive single-cell and multi-omic profiling of pleural effusions to date^7,22–24^, spanning malignant (lung, breast, and ovarian cancers) and non-malignant (heart failure) conditions. Our results establish PE not as a passive byproduct of disease, but as a dynamic and accessible biospecimen that mirrors both tumor evolution and immune response in real time. The combination of single-cell RNA sequencing, TCR repertoire profiling, and genomic analysis revealed disease-specific programs, distinct immune states, and diagnostic signals with direct clinical utility.

We showed that tumor cells in MPE transcriptionally and genomically recapitulate key features of the primary tumor, enabling non-invasive access to tumor phenotypes that would otherwise require biopsy. These tumor cells exhibit heterogeneous programs across cancer types, including EMT, apoptosis, and stemness, reflecting tissue-of-origin and metastatic adaptation. This underscores the value of MPE as a “snapshot” of tumor biology, offering a live readout of transcriptional plasticity and tumor state^25,26^.

We developed and validated a robust single-cell 50-gene MPE signature that can improve tumor cell detection over current clinical standards. Unlike traditional markers such as EPCAM, which can be downregulated in EMT or poorly differentiated tumors, our multi-gene signature achieved higher sensitivity and specificity, especially in cytology-negative cases^11,27^. This demonstrates the power of a transcriptional fingerprint over single-marker strategies, addressing a key limitation in PE diagnostics and positioning this tool for clinical deployment^28^.

From an immune perspective, MPEs revealed profound disease-specific polarization, with lung cancer PEs enriched for exhausted, metabolically rewired CD8⁺ T cells, and breast cancer PEs enriched for CD4+ helper subsets and myeloid tumor-associated cells. These patterns mirrored those in matched tumors and provided valuable insights into the immune microenvironment across cancers (Salcher et al. 2022, Wu et al. 2021 and Vázquez-García et al. 2022). TCR sequencing further enabled us to track tumor-infiltrating clones and revealed a clonal overlap between tumors and PEs. Notably, MPEs contained both expanded and memory-like CD8⁺ T cells, linking clonal architecture with the phenotypic state. The presence of OXPHOS-high exhausted clones in non-responders highlights a metabolically dysfunctional CD8⁺ compartment that may drive resistance to the therapy. This suggests that T cells in PE are not only biomarkers of immune activation but also indicators of treatment efficacy^29^. Thus, MPE-derived T cell profiles could serve as predictive tools for real-time monitoring of immunotherapy response.

Our analysis of myeloid cells further complements this view. We uncovered conserved and disease-specific tumor-associated macrophage (TAM) programs, with metabolically reprogrammed alveolar-like macrophages emerging as transcriptionally plastic and potentially immunosuppressive hubs. These findings suggest actionable myeloid targets and support the idea that PE retains the tumor-imprinted architecture of myeloid adaptation^30^. Together, our findings redefine MPE as a rich, clinically accessible biospecimen with applications that extend far beyond its current diagnostic use. Its cellular complexity, mirroring the tumor and immune landscapes, makes it a compelling candidate for use in liquid biopsy, real-time immune monitoring, and stratified therapeutic guidance. By leveraging a transcriptional signature to enhance malignancy detection, revealing T cell metabolic states predictive of treatment response, and mapping disease-specific immune programs, we established MPE as a high-resolution window into tumor–immune interactions. Future studies should prioritize longitudinal PE sampling, integration with spatial technologies, and translation of these molecular readouts into deployable clinical assays for precision oncology.

## Supplementary tables

**Supplementary table 1:**
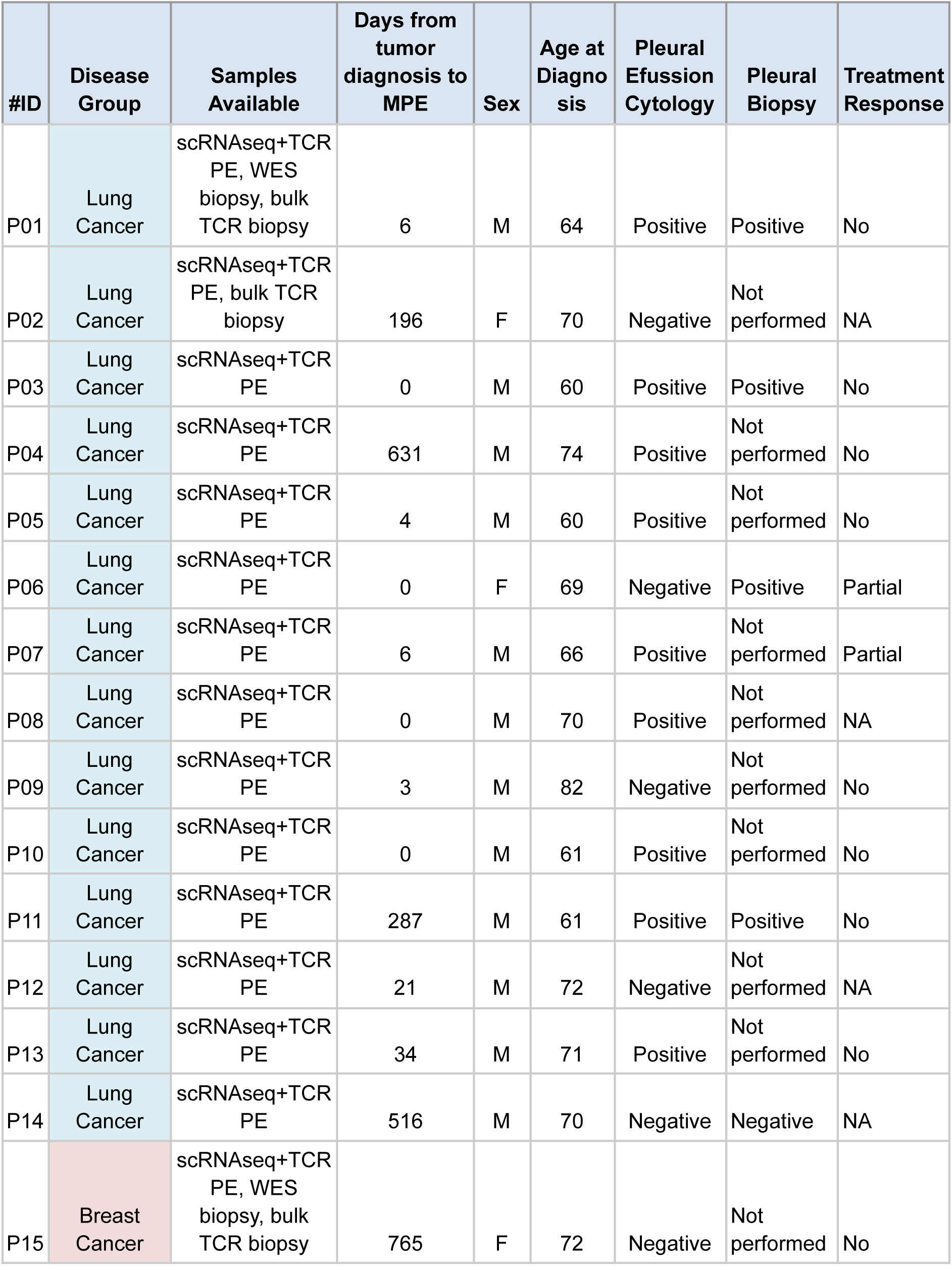
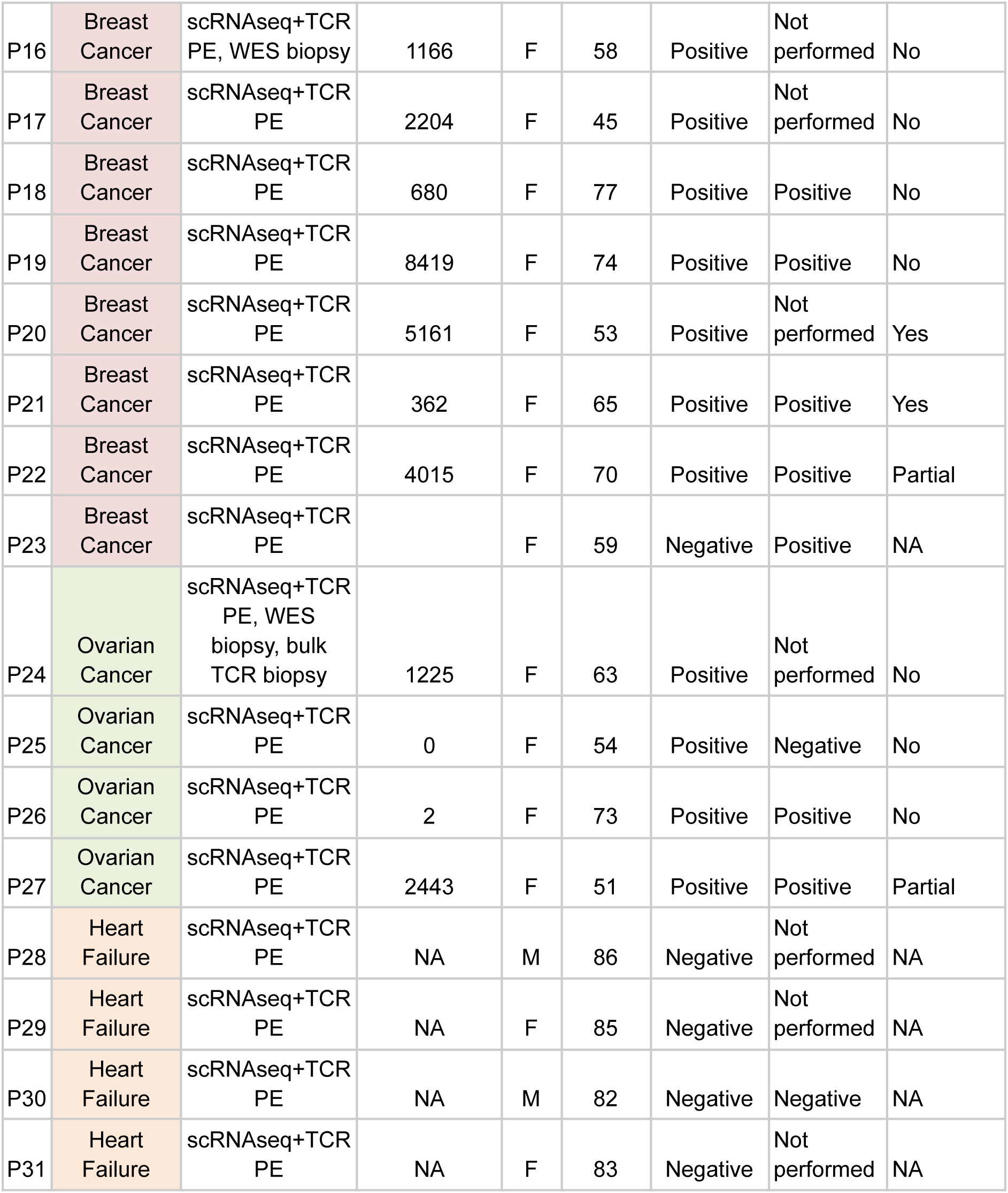
Table containing a description of the patient cohort, including clinical and demographic parameters relevant to our study.

| #ID | Disease Group | Samples Available | Days from tumor diagnosis to MPE | Sex | Age at Diagnosis | Pleural Effusion Cytology | Pleural Biopsy | Treatment Response |
| --- | --- | --- | --- | --- | --- | --- | --- | --- |
| P01 | Lung Cancer | scRNAseq+TCR PE, WES biopsy, bulk TCR biopsy | 6 | M | 64 | Positive | Positive | No |
| P02 | Lung Cancer | scRNAseq+TCR PE, bulk TCR biopsy | 196 | F | 70 | Negative | Not performed | NA |
| P03 | Lung Cancer | scRNAseq+TCR PE | 0 | M | 60 | Positive | Positive | No |
| P04 | Lung Cancer | scRNAseq+TCR PE | 631 | M | 74 | Positive | Not performed | No |
| P05 | Lung Cancer | scRNAseq+TCR PE | 4 | M | 60 | Positive | Not performed | No |
| P06 | Lung Cancer | scRNAseq+TCR PE | 0 | F | 69 | Negative | Positive | Partial |
| P07 | Lung Cancer | scRNAseq+TCR PE | 6 | M | 66 | Positive | Not performed | Partial |
| P08 | Lung Cancer | scRNAseq+TCR PE | 0 | M | 70 | Positive | Not performed | NA |
| P09 | Lung Cancer | scRNAseq+TCR PE | 3 | M | 82 | Negative | Not performed | No |
| P10 | Lung Cancer | scRNAseq+TCR PE | 0 | M | 61 | Positive | Not performed | No |
| P11 | Lung Cancer | scRNAseq+TCR PE | 287 | M | 61 | Positive | Positive | No |
| P12 | Lung Cancer | scRNAseq+TCR PE | 21 | M | 72 | Negative | Not performed | NA |
| P13 | Lung Cancer | scRNAseq+TCR PE | 34 | M | 71 | Positive | Not performed | No |
| P14 | Lung Cancer | scRNAseq+TCR PE | 516 | M | 70 | Negative | Negative | NA |
| P15 | Breast Cancer | scRNAseq+TCR PE, WES biopsy, bulk TCR biopsy | 765 | F | 72 | Negative | Not performed | No |
| P16 | Breast Cancer | scRNAseq+TCR PE, WES biopsy | 1166 | F | 58 | Positive | Not performed | No |
| P17 | Breast Cancer | scRNAseq+TCR PE | 2204 | F | 45 | Positive | Not performed | No |
| P18 | Breast Cancer | scRNAseq+TCR PE | 680 | F | 77 | Positive | Positive | No |
| P19 | Breast Cancer | scRNAseq+TCR PE | 8419 | F | 74 | Positive | Positive | No |
| P20 | Breast Cancer | scRNAseq+TCR PE | 5161 | F | 53 | Positive | Not performed | Yes |
| P21 | Breast Cancer | scRNAseq+TCR PE | 362 | F | 65 | Positive | Positive | Yes |
| P22 | Breast Cancer | scRNAseq+TCR PE | 4015 | F | 70 | Positive | Positive | Partial |
| P23 | Breast Cancer | scRNAseq+TCR PE |  | F | 59 | Negative | Positive | NA |
| P24 | Ovarian Cancer | scRNAseq+TCR PE, WES biopsy, bulk TCR biopsy | 1225 | F | 63 | Positive | Not performed | No |
| P25 | Ovarian Cancer | scRNAseq+TCR PE | 0 | F | 54 | Positive | Negative | No |
| P26 | Ovarian Cancer | scRNAseq+TCR PE | 2 | F | 73 | Positive | Positive | No |
| P27 | Ovarian Cancer | scRNAseq+TCR PE | 2443 | F | 51 | Positive | Positive | Partial |
| P28 | Heart Failure | scRNAseq+TCR PE | NA | M | 86 | Negative | Not performed | NA |
| P29 | Heart Failure | scRNAseq+TCR PE | NA | F | 85 | Negative | Not performed | NA |
| P30 | Heart Failure | scRNAseq+TCR PE | NA | M | 82 | Negative | Negative | NA |
| P31 | Heart Failure | scRNAseq+TCR PE | NA | F | 83 | Negative | Not performed | NA |

## Supplementary figures

**Supplementary Figure 1:**
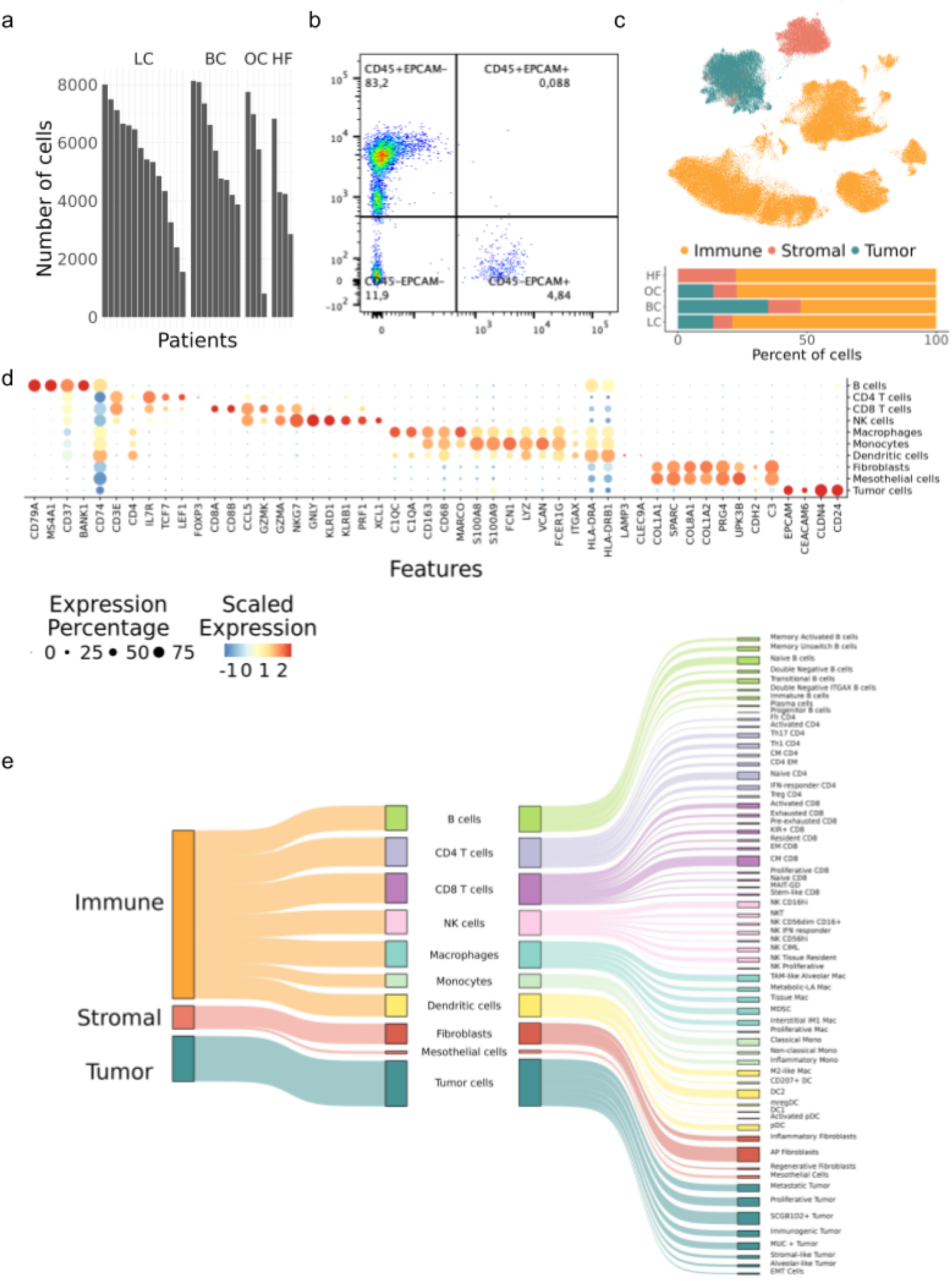
**a**, Bar plot showing the number of cells across patients and diseases. **b**, Scatter plot showing an example image obtained by flow cytometry. **c**, Integrated UMAP of all PE cells sequenced by scRNA-seq across all patients and diseases (168448 cells), colored by the general cell type (top). Bar plot showing the general cell composition across diseases (bottom). **d**, Dot plot showing the expression of marker genes across the different annotated cell types. **e**, Sankey plot showing all levels of annotation of the PE cells.

**Supplementary Figure 2:**
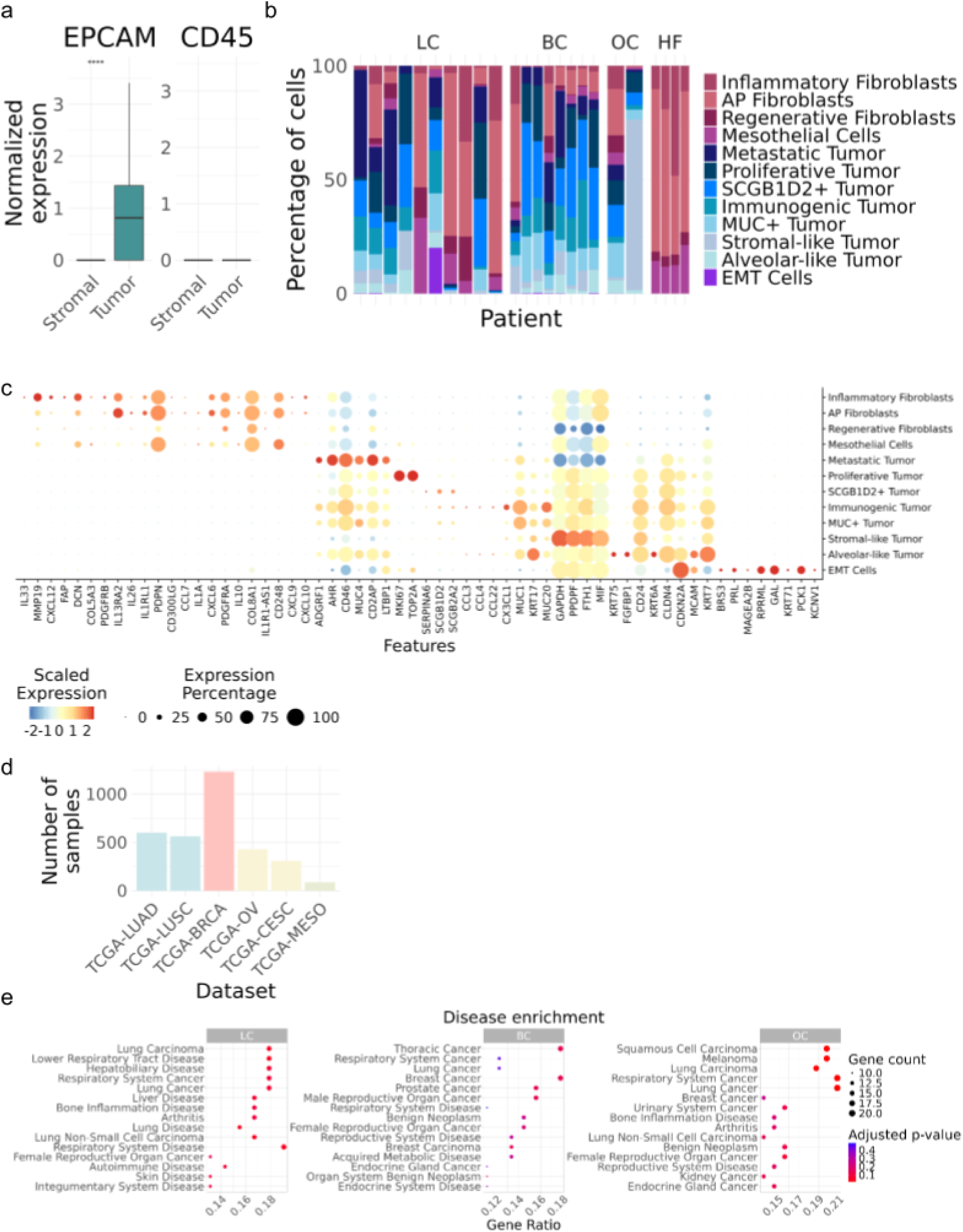
**a**, Box plot showing *EPCAM* and *CD45* gene expression distribution across stromal and tumor cell types. The Wilcoxon rank-sum test was used to compare the two groups. **b**, Bar plot showing the *CD45-* cell type composition across patients and diseases. **c**, Dot plot showing the expression of marker genes across the different annotated cell types. **d**, Bar plot showing the total number of samples included in each TCGA project (cancer type). **e**, Dot plot showing disease enrichment scores and p-values for the differentially expressed genes in each cancer type.

**Supplementary Figure 3:**
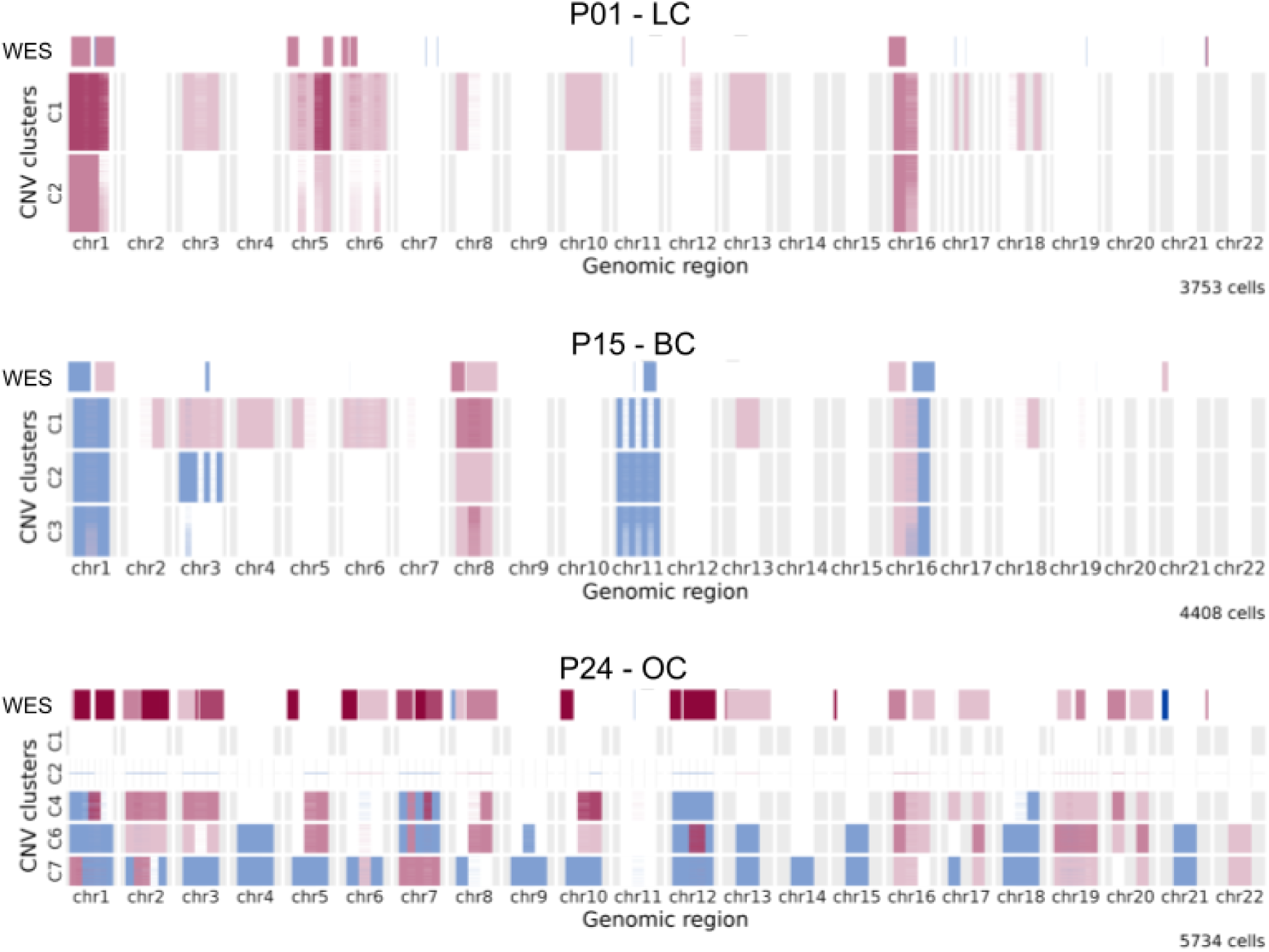
Heatmaps comparing copy number variation (CNV) profiles extracted from tumor biopsy through whole exome sequencing (WES, top line) and inferred CNV clones from the scRNA-seq tumor cells for patients for whom both samples were available.

**Supplementary Figure 4:**
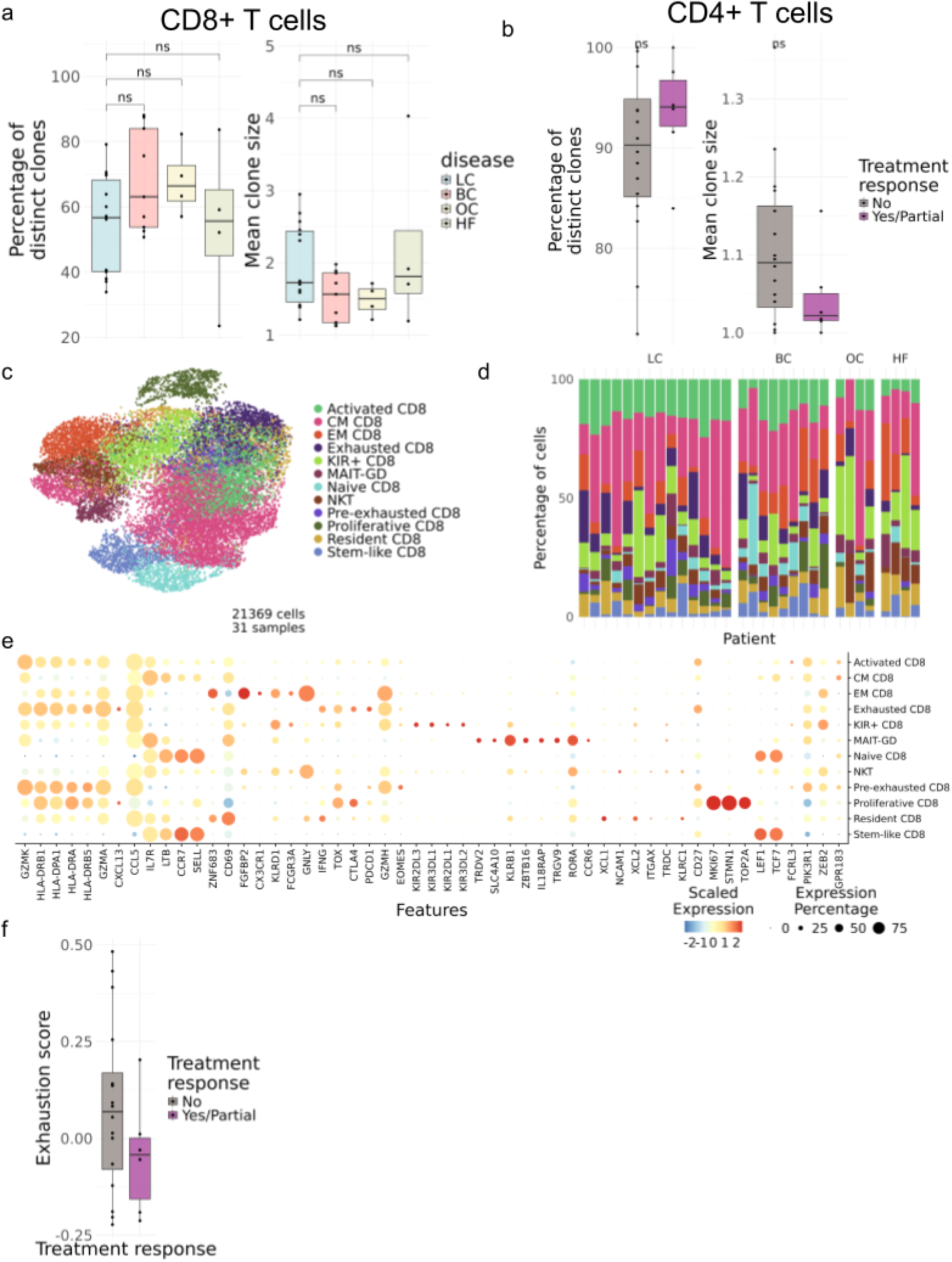
**a**, Box plots showing the percentage of distinct clones and mean clone size of CD8+ T cells per patient and disease. The Wilcoxon rank-sum test was applied to compare all disease groups with the lung cancer group. **b**, Box plots showing the percentage of distinct clones and mean clone size of*+* T cells per patient and treatment response. The Wilcoxon rank-sum test was used to compare the two groups. **c**, Integrated UMAP of all PE CD8+ T cells sequenced by scRNA-seq across all patients and diseases (21369 cells), colored by general cell type. **d**, Bar plot showing the CD8+ T cell composition across patients and diseases. Same legend as c. **e**, Dot plot showing marker gene expression across different annotated T cell types. **f**, Box plots showing the exhaustion score of CD8+ T cells per patient, grouped by treatment response.

**Supplementary Figure 5:**
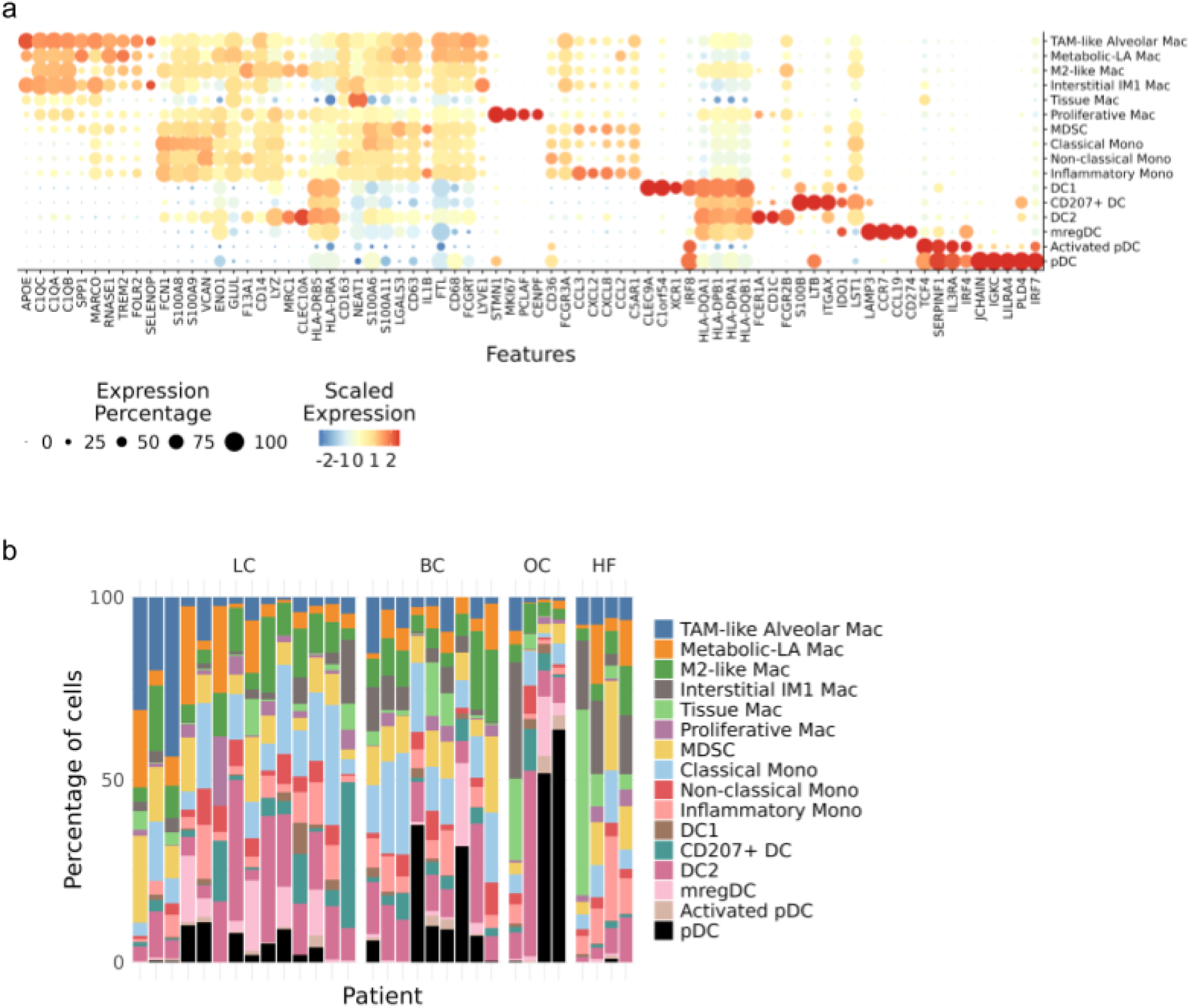
**a**, Dot plot showing marker gene expression across different annotated myeloid cell types. **b**, Bar plot showing the myeloid cell composition across patients and diseases.

## Methods

### Data generation

#### Patient cohort

The patient cohort comprised scRNA-seq data from 31 retrospectively collected PE samples. Patients with lung adenocarcinoma (n=14), breast cancer (n=9), ovarian cancer (n=4) and heart failure (n=4) PE were included. The median age at the time of PE sampling was 70 years (range, 45-86 years), with six of 19 patients being female (61.3%) and the rest, male. Additionally, five more PE samples were included for bulk RNA-seq from patients with lung adenocarcinoma (n=4) and heart failure (n=1).

Patients were diagnosed and treated at either the Hospital de la Santa Creu i Sant Pau (Barcelona, Spain) or Hospital Universitari Arnau de Vilanova (Lleida, Spain). Demographic and other clinical data (such as routine PE parameters, biopsy, and cytology status) were extracted from the patients’ records and anonymized before analysis. The clinical profiles and characteristics of the patients included in the study are summarized in **Sup. Table 1**.

#### Ethics and inclusion

This study was approved by the Institutional Ethics Committee of the Hospital de la Santa Creu i Sant Pau. Written informed consent was obtained from all patients, and samples were anonymized.

#### Collection and Processing of Pleural Fluid Samples

Pleural fluid samples were collected via thoracentesis and immediately supplemented with heparin (10 U/mL; Hospira, Lake Forest, IL, USA) to prevent clotting. Samples were processed within one hour to preserve cell viability and molecular integrity. Malignancy was determined by cytological analysis and validated by flow cytometry through detection of tumor cells. Briefly, pleural fluids were filtered through a sterile 40 µm nylon cell strainer to remove mucus and debris, and then centrifuged to obtain a cellular pellet. Red blood cells were lysed using RBC Lysis Buffer (BioLegend, San Diego, CA, USA). Subsequently, one million cells were stained with anti-CD45-PE (Immunotools, Friesoythe, Germany; clone MEM-28) and EPCAM-PECy7 (Miltenyi Biotec, Bergisch Gladbach, Germany; clone REA764) for 20 min in the dark. Cell viability was assessed using the LIVE/DEADTM Fixable Violet Dead Cell Stain Kit (Invitrogen, Carlsbad, CA, USA). Finally, the cells were washed with PBS containing 1% BSA and resuspended in 300 µL of PBS for cytometry acquisition. Doublets were excluded from the analysis. Tumor cells were identified as CD45-EPCAM+ cells.

#### Single Cell RNA-sequencing

Cryopreserved PE samples were rapidly thawed in a 37 °C water bath. Cells were transferred into a 15 ml Falcon tube containing 10 ml of pre-warmed (37 °C) RPMI medium supplemented with 10% FBS (ThermoFisher) and centrifuged at 350 × g for 7 min at room temperature (RT). The resulting cell pellet was resuspended in 1 ml of PBS supplemented with 0.05% BSA (Miltenyi Biotec), treated with DNase I (Worthington-Biochem) for 10 min at RT, and washed with an additional 9 ml of PBS/0.05% BSA. After centrifugation, the cells were counted using a TC20™ Automated Cell Counter (Bio-Rad Laboratories, S.A.). Up to 20 million cells per sample were resuspended in Cell Staining Buffer (BioLegend) and stained with antibodies for 20 min at 4 °C for fluorescence-activated cell sorting (FACS) using a Melody FACS flow cytometer (Becton Dickinson, Franklin Lakes, NJ). After staining, the cells were washed with 1 ml of staining buffer, centrifuged at 350 × g for 5 min, and resuspended in PBS/0.05% BSA. For sorting, cells were stained with anti-CD45 (PE/Cyanine7 anti-human CD45 antibody, BioLegend) and anti-EPCAM (CD326 EpCAM APC-Vio770 human, Miltenyi Biotec) antibodies to isolate the immune fraction (CD45⁺), tumor fraction (EPCAM⁺CD45⁻), and microenvironmental fraction (EPCAM⁻CD45⁻). Sorted cells were concentrated by centrifugation at 350 × g for 10 min at 4 °C and counted using a LUNA™ Dual Fluorescence Cell Counter (Logos Biosystems) after staining with Acridine Orange/Propidium Iodide. Whenever possible, the *CD45*⁺*, EPCAM*⁺*CD45*⁻, and *EPCAM*⁻*CD45*⁻ fractions were pooled in a 2:1:1 ratio prior to loading. Alternatively, the *CD45*⁺ and *CD45*⁻ fractions were directly combined in a 1:1 ratio. Pooled samples were loaded onto a Chromium X instrument (10x Genomics) for single-cell encapsulation. Maximum target cell recovery per sample was pursued, regardless of whether the Chromium Next GEM Single Cell 5’ Reagent Kit v2 (10x Genomics, PN-1000263) or GEM-X Universal 5’ Gene Expression v3 4-plex kit (10x Genomics, PN-1000780) was used. Library preparation was performed according to the manufacturer’s protocols (CG000331 and CG000770). Amplified full-length cDNAs, enriched T cell receptor (TCR) cDNAs, and final sequencing libraries were quality-checked and quantified using a high-sensitivity DNA chip on an Agilent Bioanalyzer (Agilent Technologies). Sequencing was performed on a NovaSeq 6000 system (Illumina), targeting approximately 40,000 and 10,000 read pairs per cell for gene expression (GEX) and TCR libraries, respectively. The sequencing conditions were as follows: Read 1 – 28 bp; i7 index – 10 bp; i5 index – 10 bp; Read 2 – 90 bp.

#### DNA extraction of FFPE and OCT-embedded tissue for Whole Exome and OS-TCR Sequencing

DNA was extracted from ten 3 µm-thick sections of FFPE tissue and from 5 to 10 cryosections (10 µm thick) of OCT-embedded tissue using the Maxwell® RSC FFPE Plus DNA Kit on a Maxwell® RSC instrument (Promega Corporation, Madison, WI, USA). The extracted DNA was eluted in an elution buffer containing Tris-EDTA, quantified using a NanoDrop™ ND-2000 spectrophotometer (Thermo Fisher Scientific, Waltham, MA, USA), and quality assessed based on A260/A280 ratios between 1.6 and 2.0, and A260/A230 ratios between 1.0 and 2.0. Sequencing was performed using a NovaSeq 6000 system (Illumina).

#### OS-TCR data

Extracted DNA from available samples were run with the Omniscope (Omniscope Inc., Barcelona, Spain) proprietary single-cell technology OS-T to generate sequencing libraries

### Data analysis

#### scRNA-seq data pre-processing and quality control

To profile the cellular transcriptome, we processed the sequencing reads with 10X Genomics Inc.’s software CellRanger^31^ (version 7.0.0) and mapped them against the human reference transcriptome, GRCh38 (Genome Reference Consortium Human Build 38 Organism, version 2020-A). For the TCR libraries, the corresponding VDJ human reference was also used (version 5.0.0).

We performed quality control (QC) on the raw dataset count matrices by considering the main cell metrics: number of genes, library size/number of unique molecular identifiers (UMIs) and percentage of mitochondrial RNA content per cell. Metric distributions were visualized across libraries, and consequently, we removed low-quality observations using permissive thresholds. We applied the following filters to select only high-quality cells for downstream analysis.

- Library size between 800 and 25000 UMIs
- Number of genes between 350 and 6000
- Mitochondrial content lower than 20%

The doublet detection algorithm, scrublet^32^, was applied to compute the probability of each cell barcode to capture a doublet rather than a single cell. However, we did not filter out any cells during QC based on the doublet score. During downstream analysis, we further filtered out poor-quality cell clusters that exhibited a high mitochondrial content or a high doublet score.

#### Normalization and clustering

Each scRNA-seq sample was analyzed independently to perform preliminary cell annotation, which allowed us to evaluate the integration performance in later steps. For this we used functions from the Seurat package^33^ (version 4.4.0). Normalization by library size was applied to account for differences in sequencing depth across cells. Values were then scaled by a fixed factor of 10^4 and log transformed as standard single cell best practice^34^. Following normalization, highly variable genes (HVGs) are identified to capture cell-to-cell transcriptomic variation, which is a critical step in defining cellular heterogeneity. We selected the top 3000 HVGs per sample. After HVG identification, the data were scaled to standardize the expression across genes. This centers the data by subtracting the mean expression level and dividing by the standard deviation, resulting in a z-score for each gene within each cell. Standardization is essential to ensure that differences in expression magnitudes do not disproportionately influence downstream analyses. We performed principal component analysis (PCA) and then selected the top principal components (PCs) based on the explained variance and the elbow method, to capture the most biologically relevant patterns. Subsequently, neighbor identification and clustering were applied to group cells into transcriptionally similar clusters, representing distinct cell populations. Several clustering resolutions were explored. Furthermore, Uniform Manifold Approximation and Projection (UMAP) was applied to further reduce the dimensionality of the data and visualize complex transcriptional landscapes in two dimensions. This low-dimensional visualization facilitates the exploration of cellular heterogeneity. We used the top 20 PCs for neighbor identification, clustering and UMAP.

#### Annotation

Preliminary annotation was performed by by looking at the per-cluster expression of the genes used in the sorting prior to sequencing to distinguish tumor cells (*EPCAM+*), immune cells (*PTPTC+*) and stromal cells (double negative for *EPCAM* and *PTPRC*). General annotation of the major cell populations was performed by examining the per-cluster expression of canonical genes: *CD79A, CD19* (B cells), *MZB1, IGHA1, IGHG1* (plasma cells), *CD14, CD68* (macrophages), *S100A8, LYZ, VCAN* (monocytes), *CD1C, CLEC9A, IL3R* (dendritic cells), *CD3E, CD4* (CD4+ T cells), *CD3E, CD8A, CD8B* (CD8+ T cells), *NCAM1, NKG7, GNLY* (NK cells), *UPK3B, CDH2* (mesothelial cells) and *COL1A1* (fibroblasts).

#### Integration and in-depth annotation

We then integrated the pre-processed PE samples with the python package scVI^35^ (version 1.0.4). We integrated all samples together, but also only the immune cells, myeloid cells, T cells and non-immune cells independently, to correct for technical artifacts and batch effects. For scVI integration, we adjusted the parameters (number of nodes per hidden layer, dimensionality of the latent space and number of hidden layers used for the encoder and decoder neural networks) for each dataset. Neighbor identification and clustering were performed using the python package scanpy^36^ (version 1.9.6) after integration. We followed a top-down approach, in which we first integrated and clustered all cells to annotate the major cell populations and then re-integrated and clustered each of the interesting major cell populations alone to identify subpopulations. The clustering resolution was adjusted for each dataset, and in some cases, clusters were split or merged after examining the expression profiles. In some cases, poor-quality or doublet clusters emerged and were removed from downstream analysis. UMAP was also applied after integration to obtain a harmonized two-dimensional embedding in which cells from different samples are comparable within the same low-dimensional space.

In-depth characterization of cell subpopulations was carried out by immunology and PE experts by examining the differentially expressed genes in each cluster and comparing them to populations described in the literature.

#### TCR data analysis

We defined TCR clonotypes as T cells with an exact overlap in the beta-chain receptor amino acid sequence. We used scRepertoire^37^ (version 1.12.0) to combine the gene expression and TCR information from the same cells. Only TCR sequences associated with cells annotated as T cells by RNA-seq were retained for downstream analysis.

#### Bulk RNA-seq data processing

Samples were processed using the RNA-seq pipeline (nf-core/rnaseq, v3.19.0)^38^ implemented in Nextflow version 25.04.4^39,40^.

#### WES data processing

Samples were processed using the Sarek pipeline (nf-core/sarek, v3.3.1)^41,42^ implemented in Nextflow version 23.04.3^39,40^, including copy number variation (CNV) inference through CNVkit^43^.

#### CNV analysis of scRNA-seq

Copy number profiles and clones from scRNA-seq data were inferred using CONGAS+^44^, a Bayesian framework for inferring copy number-based tumor subclones and mapping their transcriptional profiles.

#### Signature scoring and pseudobulk analysis

Signature scoring was performed in two different ways. At the single cell level, signatures were computed by cell, using the UCell R package^45^ (version 2.6.2). At the patient or disease level, we first aggregated the counts for a specific cell type or across all cells for each patient or disease by computing a “pseudobulk” using the presto R package^46^ (version 1.0.0) which was then normalized. We then scored signatures using decoupleR^47^ (version 2.8.0) on the pseudobulks by applying a multivariate linear model (MLM). The same approach was used to compute signature scores on bulk RNA-seq samples, where “pseudobulking” was not necessary, but the rest of the steps remained.

#### Compositional analysis

To estimate changes in cell population proportions across various diseases, we used the scCODA Python package^48^, a Bayesian modeling tool designed to account for the compositional nature of single-cell data and reduce the likelihood of false discoveries. This enables us to infer shifts between conditions while incorporating additional covariates. It detects differences between a reference cell type, which is assumed to remain constant across conditions, and other cell types. To conduct our analysis in an unsupervised manner, we allowed scCODA to automatically select this reference. scCODA takes as input the number of cells of each cell type in each patient and outputs the list of proportion changes for cell types, along with the corresponding corrected p-values (adjusted through the False Discovery Rate procedure, FDR).

#### Statistical analysis and data visualization

All analyses presented in this manuscript were carried out using the programming languages R versions 4.2.3 (2023-03-15) and 4.3.3 (2024-02-29), and Python versions 3.9.18 and 3.9.21. More details about the specific packages and functions used can be found in the GitHub repository associated with this publication. Detailed information on the statistical analyses and significance levels are indicated in the figure legends and text when necessary. Some illustrations were created with Biorender (https://www.biorender.com/) and all figures were assembled using Inkscape (https://inkscape.app/).

#### Box Plots

To summarize and visualize the data distribution, we used *geom_boxplot()* from the ggplot2 package in R^49^. This function generates a box plot that provides an overview of the key summary statistics, including the median, interquartile range (IQR), and potential outliers. The box plot allows for an efficient comparison of the distributions across different groups within the data. The central box represents the interquartile range (IQR) of the data, which spans from the 25th percentile (first quartile, Q1) to the 75th percentile (third quartile, Q3). Within this box, a horizontal line marks the median (50th percentile) of the data, providing a measure of the central tendency. The “whiskers” extend from the box up to a maximum of 1.5 times the IQR above Q3 and below Q1, covering most non-outlier data points. Points beyond the whiskers are considered potential outliers and are plotted as individual dots.

#### Data and code availability

Additional information is available upon reasonable request to the corresponding author.

## Declarations

### Author Contributions

**HH** and **JCN** conceived the study. **PN, HH, SV, and JCN** were involved in the study design. **HH, SV, JMP** and **JCN** supervised the project. **PN** performed all bioinformatic and statistical analyses of the data.

**MM, RO and SV** contributed to pleural fluid processing, cell isolation, and immunophenotyping analysis.

**DM** and **GC** carried out the single-cell experiments.

**JMP** was in charge of selecting the patients, obtaining pleural fluid by thoracentesis or the pleural biopsy when applicable, entering them into the database, and filling in the clinical information.

**MAS** contributed to the recruitment of patients and coordinated the IRBLleida Biobank and Anatomy Pathology service of University Hospital Arnau de Vilanova (HUAV) for biopsy retrieval, tissue sectioning, and bulk DNA extraction for downstream analyses.

**VP** and **AP** contributed to patient recruitment, data collection, and the development and maintenance of the study database.

**AR** and **PM** contributed to patient recruitment and data collection.

**PN** and **JCN** interpreted the results and drafted the manuscript. All authors contributed to the critical discussion of the manuscript and read and approved the final version.

## Acknowledgements

We thank the patients and their families, without whom this research would not have been possible. We acknowledge the TCGA Research Network, whose data helped validate our work.

